# A metabolic crosstalk between liposarcoma and muscle sustains tumor growth

**DOI:** 10.1101/2023.06.02.543362

**Authors:** Manteaux Gabrielle, Prieto Romero Jaime, Gayte Laurie, Riquier-Morcant Blanche, Amsel Alix, Jacq Solenn, Cisse Madi Y, Perrot Gaelle, Chibon Frédéric, Pomies Pascal, Carrere Sebastien, Firmin Nelly, Riscal Romain, Linares Laetitia K

## Abstract

Dedifferentiated (DD-LPS) and Well-differentiated (WD-LPS) liposarcoma are characterized by a systematic amplification of the *MDM2* oncogene. We recently demonstrated that p53-independent metabolic functions of chromatin-bound MDM2 (C-MDM2) are exacerbated in LPS and mediate an addiction to serine metabolism in order to sustain tumor growth. Here, we show that metabolic cooperation between LPS and distant muscle, which raise serine and glycine blood levels, is essential for LPS tumor growth. By releasing IL-6, tumor influence distant muscle to upregulate their serine synthesis machinery. Blocking IL-6 secretion or treating LPS cells with FDA approved IL-6 inhibitor, decreased serine production and impaired tumor proliferation. These data reveal IL-6 as a central tumorkine in metabolic crosstalk between tissues and identifies IL-6 as a plausible treatment for LPS patients.

## INTRODUCTION

Sarcomas, which represent about 1% of all cancers, are malignant tumors of mesenchymal origin arising from soft or bone tissues. Among the 100 different histological subtypes of sarcomas, liposarcoma (LPS) represent the second most frequent subtype after gastrointestinal stromal tumors (GIST), accounting for 15-20% of all sarcomas^1^. The prognosis of LPS is very heterogeneous and depends on the tumor location, its histological subtype, and the grade/size of the tumor at diagnosis^2–4^. The risk of recurrence and metastatic dissemination of advanced LPS varies between 20 to 40% in case of localized tumors, and mainly depends on the quality of their surgical resection that remains the most efficient therapeutic strategy to date. LPS are known to be poorly responsive to classical chemotherapies and despite that new targeted therapies, such as tyrosine kinase inhibitor Pazopanib, have demonstrated efficacy in patients with several types of sarcomas, they showed no benefit to patients with LPS^5^. The median overall survival estimated around 15 months^6, 7^, and the absence of treatment for metastatic or unresectable LPS reveal an urge for novel therapeutic strategy for LPS.

The most common LPS subtypes, the well differentiated and the dedifferentiated LPS (WD-LPS and DD-LPS, respectively), are characterized by the systematic amplification of the q13-15 region of chromosome 12. This region contains the "*murine double-minute 2*" gene (*Mdm2*), which encodes a well-characterized negative regulator of the p53 tumor suppressor. The frequency of *Mdm2* amplification is such (almost 100%) that it is currently used for routine diagnosis to distinguish WD/DD-LPS from other sarcoma subtypes that commonly harbor p53 mutations^8, 9^.

The *mdm2* gene was first identified as the gene involved in the spontaneous transformation of an immortalized murine cell line, BALB/c 3T3. *Mdm2* was later labeled as an oncogene participating in cell transformation. MDM2 oncoprotein is frequently overexpressed in numerous human cancers ^10^, resulting in a loss of the tumor suppressor p53-dependent activities^11^. Under normal growth conditions, the p53 protein is kept at low levels by MDM2-mediated polyubiquitylation inducing its degradation by the 26S proteasome. The role of MDM2 as a major regulator of p53 stability and transcriptional activities has been widely described by *in vitro* and *in vivo* models^11–13^.

Through its E3 ligase activity, MDM2 affects the function of other cellular proteins involved in cell proliferation, DNA repair, ribosome biosynthesis, and many other processes that can also contribute to its oncogenic potential^14, 15^. However, growing evidences suggest that MDM2 is involved in a complex network of protein interactions that confer MDM2 new functions beyond its relationship with p53^16–19^. More recently, we started an in-depth analysis of MDM2 functions independently of p53 and demonstrated a key role in serine metabolism^20^.

We have shown that MDM2 is recruited on chromatin (C-MDM2) in a p53 independent manner and regulates an ATF-3 and ATF-4 dependent transcriptional program. Although we described that MDM2 is mainly recruited on chromatin under specific stress conditions such as, oxidative stress. Moreover, performing a screen on a large panel of cancer cell lines, we identify liposarcoma cell lines as the one which consistently and spontaneously harbor C-MDM2^20^. The p53-independent metabolic functions of C-MDM2 are exacerbated in LPS and mediate an addiction to serine metabolism that sustains nucleotide synthesis and tumor growth^21^. Treatment of LPS cells with Nutlin-3A, a pharmacological inhibitor of the MDM2-p53 interaction, stabilizes p53 but unexpectedly enhances MDM2-mediated control of serine metabolism by increasing its recruitment to chromatin, likely explaining the poor clinical efficacy of this MDM2 inhibitors class^22^. In contrast, genetic or pharmacological inhibition of C-MDM2 by SP141, a distinct MDM2 inhibitor triggering its degradation, and interfering with *de novo* serine synthesis, impaired LPS growth both *in vitro* and in clinically relevant patient-derived xenograft models. Taken together, our data suggest that targeting MDM2 functions in serine metabolism represents a potential therapeutic strategy for LPS^21, 23^.

The impact of cancer cells on their environment, locally and distantly, is known to promote malignancy and chemoresistance ^24^. Understanding the interactions between cancer cell and surrounding metabolism will be critical for combining metabolism-targeted therapies with existing chemotherapies. It is known that cancer cells compete with cellular components of the microenvironment for essential nutrients, such as glucose, amino acids and lipids. For example, restricting T cell glucose metabolism causes lymphocyte exhaustion^25^, when high arginine levels promotes enhanced T cell survival and anti-tumor activity^26, 27^. The relationship between tumors and other organs is not limited to the immune system. Endothelial cells also undergo factor-induced metabolic reprogramming^28^. Cancer also causes alterations in whole-body metabolism that may influence how tumor access to essential resources. Acting on nutrient availability through diet composition has been shown to slow cancer progression^29^ and this field is likely to be a productive area of research in the near future.

In this study, we demonstrate a new concept in liposarcoma pathogenesis, related to a potential metabolic “long-distance” cooperation between normal tissues and LPS. Our data obtained in humanized patient-derived mouse models of LPS (PDX) suggests that LPS somehow "educate" normal tissues/cells to support their massive demand in serine. While monitoring LPS growth *in vivo*, we observed an increase of circulating serine and glycine levels. Interestingly, skeletal muscles of these engrafted animals were found to upregulate a transcriptional program, involved in *de novo* serine synthesis and described to be regulated by C-MDM2 in cancer cells^21^. We hypothesized that a metabolic cooperation between LPS cancer cells and surrounding muscle allows LPS tumors to maintain serine pools and now identify interleukin-6 (IL-6), as essential for LPS mediated muscle reprogramming. Furthermore, blocking IL-6 using an antibody, results in exogenous serine starvation in LPS cells and impaired proliferation *in vitro* and *in vivo*. As such, the exogenous serine provided by surrounding tissue, such as muscle, is critical for LPS tumor growth and reveals IL-6 as a plausible target for novel LPS treatments.

## RESULTS

### Muscle reprogramming fuels Liposarcoma serine addiction

Recently, we classified liposarcoma as serine-dependent tumors that use both, self-serine production and serine auxotrophy ^21^. To further investigate serine metabolism significance in liposarcoma, we looked at circulating serine and glycine level in nude mice xenografted with LPS cell lines or humanized patient-derived mouse models (PDX). PDX are patient-derived tumor xenograft models generated upon transfer of freshly resected human tumor samples of primary naive LPS into immunodeficient nude mice. Surprisingly, we observed that serine and glycine levels in the blood stream are significantly higher in mice harboring LPS tumor relative to control mice. Additionally, in response to MDM2 inhibitor treatment, which induces a drastic reduction in tumor growth ^21^, we were able to rescue serine and glycine levels at the base line observed pre engraftment, confirming a correlation between circulating levels of serine and glycine and LPS tumorigenesis (Fig. 1A-D). Given that more than 90% of the serine consumed by tumor cells comes from exogenous sources^21^, we anticipated that liposarcoma tumors depend on serine and glycine synthetized by surrounding tissue. To test this hypothesis, we examined the expression of genes encoding limiting enzymes of the *de novo* serine synthesis pathway (SSP), including *3-phosphoglycerate dehydrogenase* (*Phgdh*), *phosphoserine aminotransferase 1* (*Psat1*), and *phosphoserine phosphatase* (*Psph*) in distant metabolic organs, such as liver and muscles. Interestingly, these genes were upregulated in skeletal muscles of engrafted animals (Fig. 1E). In contrast, liver, kidney, brain and adipose tissue did not exhibit such activation of SSP genes (Fig. 1F and Extended Data Fig. 1A-C). These results indicate that muscle reprogramming, and no other tissues, sustains liposarcoma need in serine and glycine amino acids. To further investigate the specificity of muscle-secreted serine on liposarcoma tumorigenesis, control mice fed with serine/glycine-deprived diet were analyzed. As expected, serine and glycine levels in blood are lower (Extended Data Fig. 1D) and because liver and kidney are known to be essential organs involved in serine/glycine homeostasis^30^, SSP genes were activated in these tissues but not in muscle, brain or adipose tissue (Fig. 1G, H and Extended Data Fig. 1E-G) confirming the specificity of this muscle reprogramming observed in mice harboring LPS. Of note, in mice with C-MDM2 independent breast cancer PDXs, the *de novo* serine synthesis transcriptional program regulated by C-MDM2 was not activated (Fig. 1I). Collectively, these results indicate that liposarcoma tumors potentially induce skeletal muscle reprogramming to sustain serine levels required for proliferation and survival.

**Figure 1:**
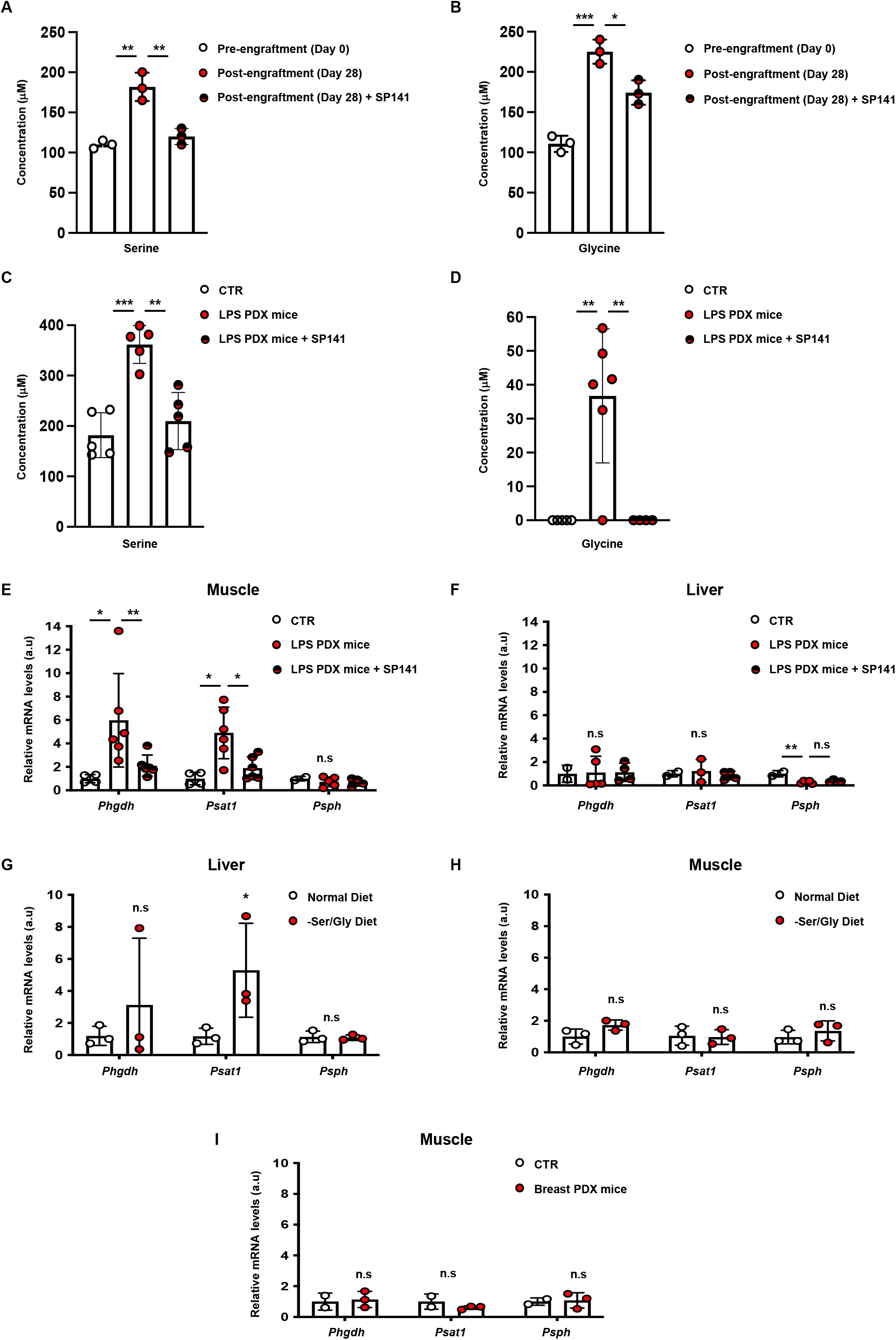
Liposarcoma reprogram serine synthesis pathway in muscles. **A**, Serine and **B**, Glycine levels (μM) measured by HPLC, in nude mice before and 28 days post-engraftment of liposarcoma cells (IB115). Mice were treated daily with placebo or SP141 (40mg/ml). **C**, Serine and **D**, Glycine levels (μM) measured by HPLC. Control mice were compared to liposarcoma PDX Mice (LPS-PDX). Mice were treated daily with placebo or SP141 (40mg/ml). **E**, Real-time qPCR analysis performed on LPS-PDX and control mice muscle, evaluating expression of serine synthesis pathway genes: *Phgdh, Psat1* and *Psph*. Mice were treated daily with placebo or SP141 (40mg/ml). **F**, Real-time qPCR analysis performed on LPS-PDX and control mice liver, evaluating expression of serine synthesis pathway genes: *Phgdh, Psat1* and *Psph*. Mice were treated daily with placebo or SP141 (40mg/ml). **G**, Real-time qPCR analysis performed control mice liver, evaluating expression of serine synthesis pathway genes: *Phgdh, Psat1* and *Psph*. Mice were fed with normal or -Ser/Gly diet. **H**, Real-time qPCR analysis performed on control mice muscle, evaluating expression of serine synthesis pathway genes: *Phgdh, Psat1* and *Psph*. Mice were fed with normal or -Ser/Gly diet. **I**, Real-time qPCR analysis performed on Breast-PDX and control mice muscle, evaluating expression of serine synthesis pathway genes: *Phgdh, Psat1* and *Psph*. (All experiments were performed in at least triplicates and statistical analysis was applied with *=P<0.05, **=P<0.01, ***=<0.001, n.s=non-significant).

### Liposarcoma cells take advantage of surrounding muscle cells for growth

To understand mechanisms by which liposarcoma cancer cells reprogram muscle cells, mRNA levels of SSP enzymes were assessed in co-cultures of human liposarcoma cells IB115 or IB111 and murine myoblast C2C12. It should be noted that direct interaction between human and murine cells allowed us to discriminate genes expression from the two population of cells present in the same dish using specific primers. Interestingly, we demonstrated that C2C12 co-cultured with LPS cells exhibit an increase of SSP genes expression compared to C2C12 cultured alone or co-cultured with other cancer type, such as breast cancer (MCF7) and melanoma (A375) cell lines (Fig. 2A and Extended Data Fig. 2A-C). Moreover, because we observed a distant muscle reprogramming in mice, we suggest that liposarcoma tumors could produce factors invoking muscle metabolism rewiring. To decipher these mechanisms, we collected culture media from LPS cells; conditioned media (CM), and test the impact of CM on C2C12 SSP genes. CM from IB115 and IB111 were added to C2C12 cells for 48h and mRNA extraction was used to evaluate SSP genes response. Interestingly, CM from LPS cancer cell lines increased *Psat1, Phgdh and Psph* mRNA expression in C2C12 indicating that a factor released specifically by liposarcoma cells induces serine metabolism reprogramming in muscle cells (Fig. 2B, C). Similar results were obtained using myotubes and luciferase technology (Extended Data Fig. 2D, E). We also confirmed these results using transformed human myoblast (Myo-E7) (Extended Data Fig. 2F). To guarantee the exclusiveness of the special crosstalk between liposarcoma and muscle cells, liposarcoma cells were co-cultured with murine liver cells, BMEL, and we observed that SSP genes were not induced (Fig. 2D) Previous report from our lab demonstrated that ATF4 dependent C-MDM2 regulates the transcription of serine metabolism genes in LPS^21^. To assess the role of Mdm2 in our model, we targeted Mdm2 in C2C12 cells (C2C12 sh*Mdm2*). Whereas IB115 induced the 3 SSP genes in C2C12 sh*Scr*, IB115 failed to induce the same transcriptional program in sh*Mdm2* expressing C2C12 (Fig. 2E, F and Extended Data Fig. 2G, H). Depleting Mdm2 co-transcription factor, Atf4 by shRNA in C2C12 also lead to similar results (Fig. 2G, H and Extended Data Fig. 2I, J) confirming the importance of Mdm2-Atf4 complex in the activation of the SSP transcriptional program LPS-mediated in C2C12. To confirm that increase serine *de novo* biosynthesis in the myoblast is associated with intercellular serine releasing, we assessed serine levels in C2C12 media. As expected, serine levels were higher in C2C12 media cocultured with IB115 compared to IB115 alone (Fig. 2I).

**Figure 2:**
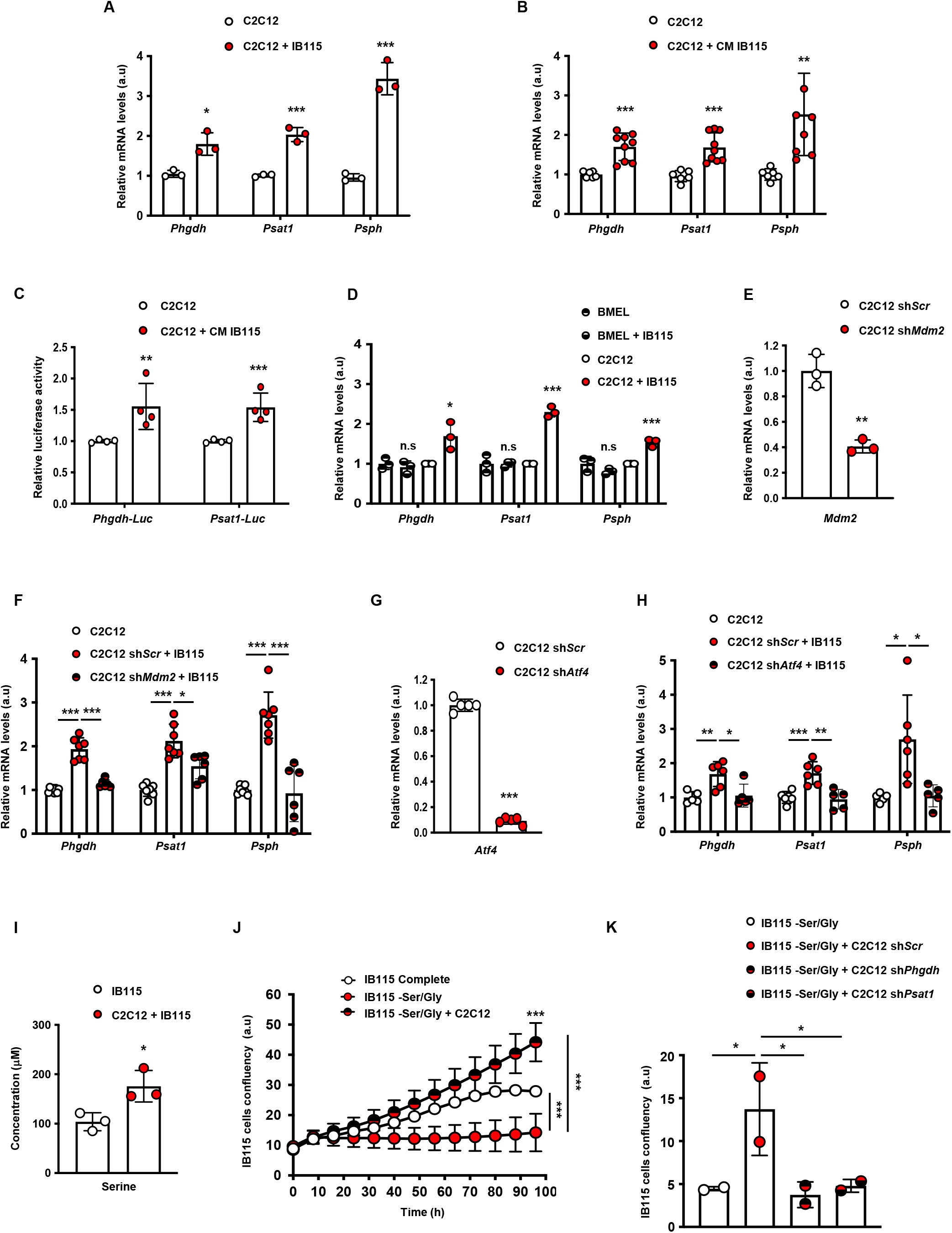
Liposarcoma needs muscle reprogramming to sustain proliferation. **A**, Real-time qPCR analysis performed on C2C12 cocultured with IB115 cells (ratio 1:2), evaluating expression of serine synthesis pathway genes: *Phgdh, Psat1* and *Psph*. **B**, Real-time qPCR analysis performed on C2C12 cells incubated 16h with IB115 conditioned media, evaluating expression of serine synthesis pathway genes: *Phgdh, Psat1* and *Psph*. **C**, Luciferase assay performed on C2C12 cells incubated 16h with IB115 conditioned media, evaluating relative luciferase activity of *Phgdh* and *Psat1* reporter. **D**, Real-time qPCR analysis performed on C2C12 or BMEL murine cells cocultured with IB115 cells (ratio 1:2), evaluating expression of serine synthesis pathway genes: *Phgdh, Psat1* and *Psph*. **E**, Real-time qPCR analysis of *Mdm2* mRNA level in C2C12 cells after sh*Scr* or sh*Mdm2* lentiviral infection, puromycin selection (48h, 2μg/mL). **F**, Real-time qPCR analysis performed on C2C12 cells after sh*Scr* or sh*Mdm2* cocultured with IB115 cells (ratio 1:2), evaluating expression of serine synthesis pathway genes: *Phgdh, Psat1* and *Psph*. **G**, Real-time qPCR analysis of *Atf4* mRNA level in C2C12 cells after sh*Scr* or sh*Atf4* lentiviral infection, puromycin selection (48h, 2μg/mL). **H**, Real-time qPCR analysis performed on C2C12 cells after sh*Scr* or sh*Atf4* cocultured with IB115 cells (ratio 1:2), evaluating expression of serine synthesis pathway genes: *Phgdh, Psat1* and *Psph*. **I**, Serine levels (μM) measured by HPLC in IB115 alone or cocultured with C2C12 cells (ratio 1:2). **J**, Proliferation assay performed on IB115 cells grown in media supplemented with or without Serine and Glycine and cocultured with C2C12 cells. **K,** End point of proliferation assay performed on IB115 cells grown in media without Serine and Glycine and cocultured with C2C12 cells after sh*Scr*, sh*Phgdh* or sh*Psat1* cells. (All experiments were performed in at least triplicates and statistical analysis was applied with *=P<0.05, **=P<0.01, ***=<0.001, n.s=non-significant).

Given that liposarcoma induces muscle reprogramming, we anticipated that they depend on muscle serine production for growth. To test this hypothesis, LPS were cultured in media without serine and glycine (-Ser/Gly) and in the presence or not of C2C12 cells. We observed proliferation defect in the absence of serine and glycine relative to cells grown in full media (Complete). Interestingly, co-culturing LPS cells with C2C12 in -Ser/Gly, fully rescued LPS cells proliferation (Fig. 2J). Similar results were obtained using human myoblast (Extended Data Fig. 2K). In contrast, co-culturing LPS cells with C2C12 lacking enzymes involved in the SSP (C2C12 sh*Psat1*, sh*Phgdh*) failed to rescue IB115 proliferation (Fig. 2K and Extended Data Fig. 2L, M). Collectively, these results indicate that LPS cells require muscle cells to reach serine requirement and sustain proliferation in a MDM2 dependent manner.

### Liposarcoma-released IL-6 regulates muscle reprogramming

We hypothesized that soluble factors might be released by LPS cells to initiate serine metabolism in distant muscle. To test the extent to which secreted factors from the liposarcoma cells promote muscle reprogramming, we collected LPS cells media. The analysis of this culture supernatant using a cytokines array shows that LPS cells release significant number of different cytokines including interleukin-6 (IL-6) (Fig. 3A). Interestingly, previous reports linked ATF4 with IL-6 expression in macrophages^31, 32^. Furthermore, MDM2 has been involved in a IL-6-mediated degradation of p53^33^. For those reasons, we decided to concentrate our effort on IL-6. We then examined *IL-6* gene expression in liposarcoma cell lines relative to other cancer cell lines and observed higher *IL-6* mRNA levels in LPS cell lines (Fig. 3B). IL-6 protein levels were also overrepresented in supernatant of liposarcoma cell lines analyzed by ELISA (Fig. 3C). In addition, to confirm the efficiency of IB115-released IL-6, we treated IL-6-dependent myeloma cell line XG-6 with IB115 CM. CM from IB115 was able to increase XG-6 proliferation while CM from MCF7 not (Fig. 3D and Extended Fig. 3A). To further investigate by which mechanism IL-6 is released from LPS tumors, we performed ChIP experiments and observed ATF4 and MDM2 binding on *IL-6* promoter (Fig. 3E), suggesting that ATF4/MDM2 complex regulates *IL-6* transcription. Moreover, we confirmed IL-6 regulation through ATF4 and MDM2 in LPS cells using shRNA technology. As expected, *IL-6* mRNA in LPS cells and excreted IL-6 protein from LPS cells were lowered upon shRNA-*ATF4* and –*MDM2* treatment (Fig. 3F, G and Extended Fig. 3B, C).

**Figure 3:**
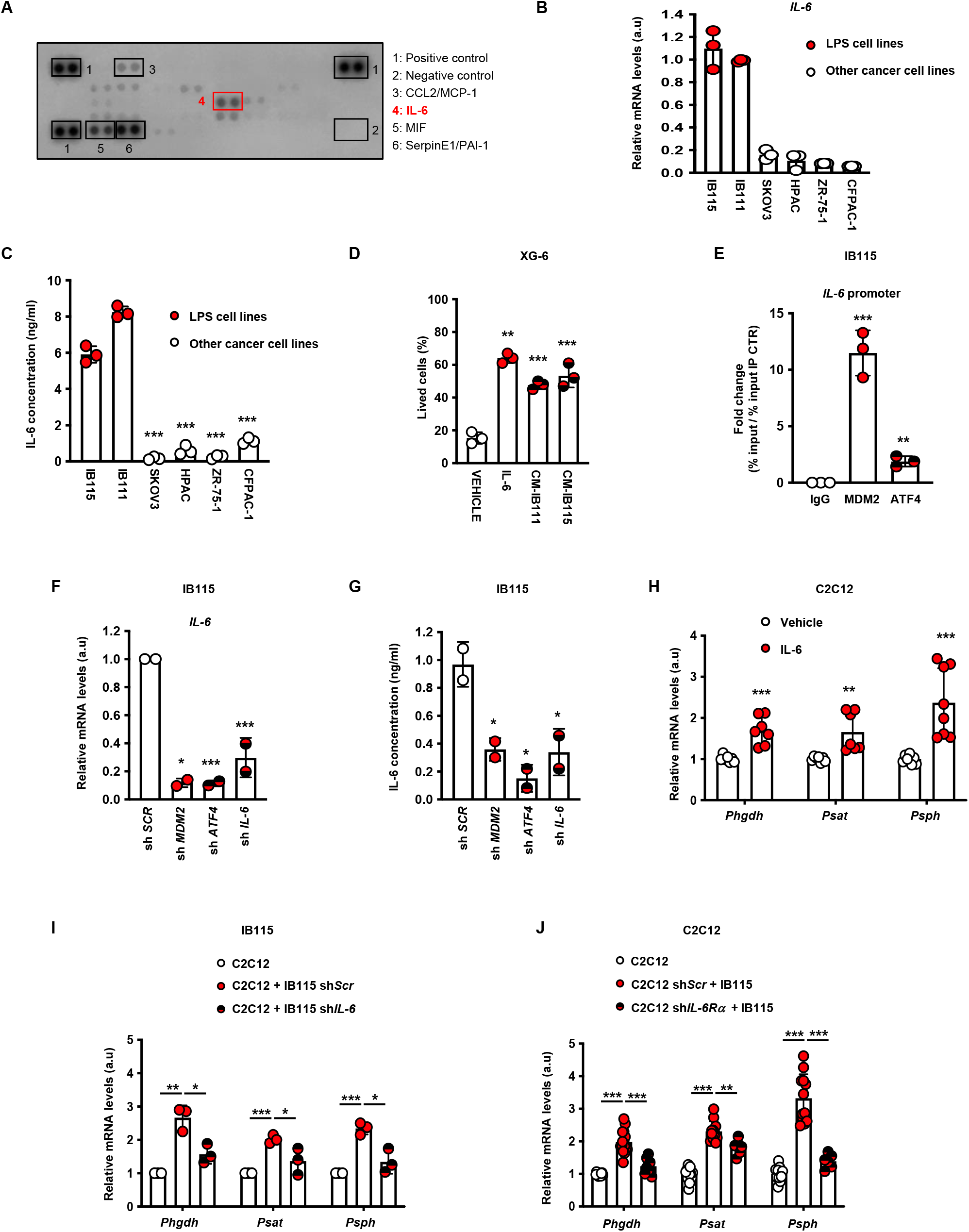
Liposarcoma-released IL-6 reprograms distant muscles **A,** Proteome Profiler Human Cytokine Array. **B,** Real-time qPCR analysis of *IL-6* mRNA level of different cancer cell lines, SKOV3, HPAC, ZR-75.1, CFPAC-1, including LPS cell lines, IB115 and IB111. **C,** IL6 content (ng/ml), measured by Elisa, of different cancer cell lines, SKOV3, HPAC, ZR-75.1, CFPAC-1, including LPS cell lines, IB115 and IB111. **D,** End point of proliferation assay performed on XG-6 cells grown in media supplemented with vehicle, recombinant IL-6, IB115 or IB111 conditioned media. **E,** Chromatin immunoprecipitation-qPCR experiment performed IB115 cells. Results are represented as the relative ratio between the mean value of immunoprecipitated chromatin (calculated as a percentage of the input) with the indicated antibodies. **F,** Real-time qPCR analysis of *IL-6* mRNA level in IB115 cells after sh*Scr*, sh*Mdm2,* sh*Atf4 or* sh*IL-6* lentiviral infection, puromycin selection (48h, 2μg/mL). **G,** IL6 content (ng/ml), measured by Elisa, of IB115 cells after sh*Scr*, sh*Mdm2,* sh*Atf4 or* sh*IL-6* lentiviral infection, puromycin selection (48h, 2μg/mL). **H,** Real-time qPCR analysis performed on C2C12 cells grown in media supplemented with vehicle or recombinant IL6, evaluating expression of serine synthesis pathway genes: *Phgdh, Psat1* and *Psph*. **I,** Real-time qPCR analysis performed on IB115 cells after sh*Scr* or sh*IL-6* cocultured with C2C12 cells (ratio 1:2), evaluating expression of serine synthesis pathway genes: *Phgdh, Psat1* and *Psph*. **J,** Real-time qPCR analysis performed on C2C12 cells after sh*Scr* or sh*IL-6Ra* cocultured with IB115 cells (ratio 1:2), evaluating expression of serine synthesis pathway genes: *Phgdh, Psat1* and *Psph*. (All experiments were performed in at least triplicates and statistical analysis was applied with *=P<0.05, **=P<0.01, ***=<0.001, n.s=non-significant).

To determine the uncharacterized role of IL-6 on muscle reprogramming, we analyzed the level of SSP genes in C2C12 murine myoblast cultured in the presence of recombinant IL-6. Interestingly, 50 pg/ml of IL-6 were sufficient to activate serine synthesis genes expression in C2C12 (Fig. 3H) We also assess the effect of targeting the IL-6 pathway operating between LPS cells and distant muscle. Serine genes activation was not observed in C2C12 cells when cocultured with IB115 sh*IL-6* (Fig. 3I and Extended Data Fig. 3D, E). Similar results were observed with C2C12 sh*IL-6R* cocultured with IB115 (Fig. 3J and Extended Data Fig. 3F, G). Collectively, these results indicate that liposarcoma cells-released IL-6 promotes serine metabolism reprogramming on muscle cells.

### Targeting the IL-6/STAT3 pathway impairs serine biosynthesis activation in reprogrammed muscle cells

Being a cytokine with numerous functions, IL-6 affects metabolism of many organs, such as white adipose tissue, liver, skeletal muscle, or pancreas ^34^. The main approach for inhibition of IL-6-mediated signaling is currently the use of antibodies. LPS serine-deprived cell lines treated with three different IL-6 antibodies (BE8, Siltuximab, hBE8) exhibited proliferation defects reflecting the absence of IL6-mediated *de novo* serine induction in C2C12 cells (Fig. 4A, B and Extended Data Fig. 4A-D). Similar results were observed with bazedoxifene (BZA), drug-targeting GP130 (IL-6Ra) (Fig. 4C and Extended Data Fig. 4E). Mechanistically, to activate its classic signaling pathway, IL-6 binds to its membrane-bound receptor (IL-6Ra) inducing the JAK (Janus Kinase) activation cascade, which then serve as docking sites for proteins initiation such as, PI3K/AKT, MAPK, Ras or STAT3 signaling pathways^35^. Our work and other have shown a link between ATF4/MDM2^20^ and ATF4/STAT3^36^ suggesting that JAK/STAT3 signaling pathway could play a prominent role in mediating effects of IL-6 on muscle reprogramming. We hypothesized that our observed IL-6-dependent proliferation phenotype was at least partially mediated through STAT3 signaling axis in C2C12. STAT3 phosphorylation was induced in C2C12 myoblasts stimulated by IB115 CM and correlate with accumulation of the 3 SSP enzymes (Fig.4D, E). Moreover, STAT3 inhibitors also led to serine metabolism transcriptional program and LPS proliferation defects (Fig.4F, G and Extended Data Fig. 4F). Therefore, how STAT3 is connected to serine biosynthesis will required more investigation in the future. Taken together, our data suggest that LPS-released IL-6 appears to be critical to sustain STAT3 signaling and control muscle reprogramming. Blocking LPS-muscle crosstalk through IL-6 inhibitors and/or serine restriction could offer a potential combination therapy for patients with liposarcoma.

**Figure 4:**
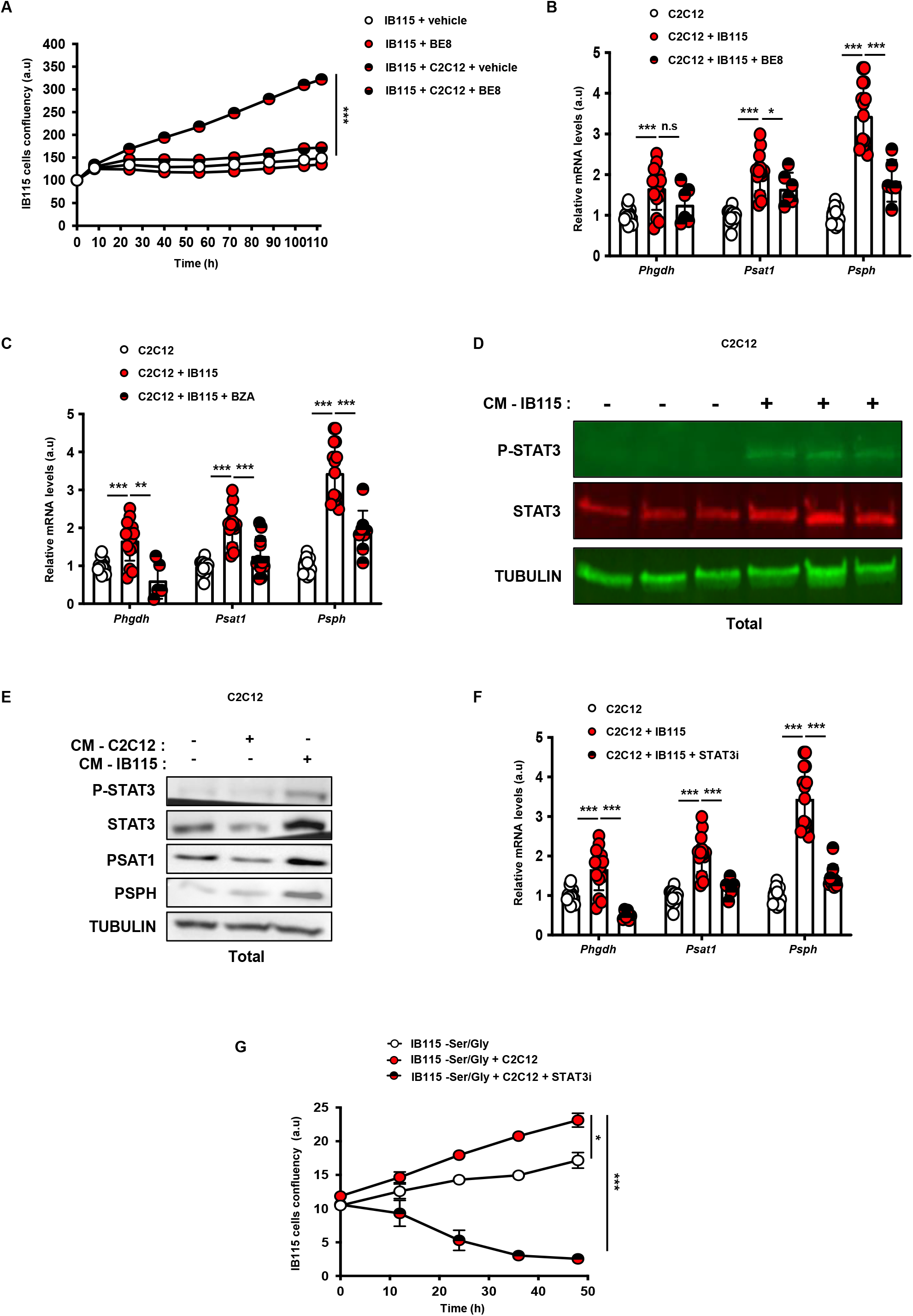
Targeting IL-6/STAT3 pathway impairs liposarcoma proliferation **A,** Proliferation assay performed on IB115 cells alone or cocultured with C2C12 cells grown in media without Serine and Glycine and supplemented with vehicle or anti IL-6 (BE8). **B,** Real-time qPCR analysis performed on C2C12 cells alone or cocultured with IB115 cells grown in media supplemented with vehicle or anti IL-6 (BE8), evaluating expression of serine synthesis pathway genes: *Phgdh, Psat1* and *Psph*. **C,** Real-time qPCR analysis performed on C2C12 cells alone or cocultured with IB115 cells grown in media supplemented with vehicle or GP130 inhibitor (BZA), evaluating expression of serine synthesis pathway genes: *Phgdh, Psat1* and *Psph*. **D,** STAT3 and P-STAT3 protein expression assessed by fluo-immunoblots in C2C12 cells incubated 16h with IB115 conditioned media. TUBULIN was used as the loading control. **E,** STAT3, P-STAT3, PSAT1 and PSPH protein expression assessed by immunoblots in C2C12 cells incubated 16h with IB115 or C2C12 conditioned media. TUBULIN was used as the loading control. **F,** Real-time qPCR analysis performed on C2C12 cells alone or cocultured with IB115 cells (ratio 1:2) grown in media supplemented with vehicle or STAT3 inhibitor, evaluating expression of serine synthesis pathway genes: *Phgdh, Psat1* and *Psph*. **G,** Proliferation assay performed on IB115 cells alone or cocultured with C2C12 cells grown in media without Serine and Glycine and supplemented with vehicle or STAT3i. (All experiments were performed in at least triplicates and statistical analysis was applied with *=P<0.05, **=P<0.01, ***=<0.001, n.s=non-significant).

### IL-6 as a novel target in liposarcoma

After the discovery of IL-6/STAT3/Serine axis and its role in liposarcoma carcinogenesis, we proposed that immunotherapy using IL-6 antibody could be useful to reduce liposarcoma tumor proliferation on mice and becoming the mainstay for future liposarcoma diagnosis and treatment. To evaluate the clinical relevance of our findings, we first measured circulating IL-6 levels in mice harboring LPS-PDX relative to normal mice. Elisa analysis revealed a higher IL-6 blood concentration in the presence of liposarcoma (Fig. 5A). Moreover, we compared *IL-6* mRNA expression in human tumor samples from different type of cancers. As expected, liposarcoma samples exhibit elevated *IL-6* mRNA abundance compared to other cancer types (Fig. 5B). Collectively, these results suggest that IL-6 screening could be potentially used to detect liposarcoma before any clinical signs appear.

**Figure 5:**
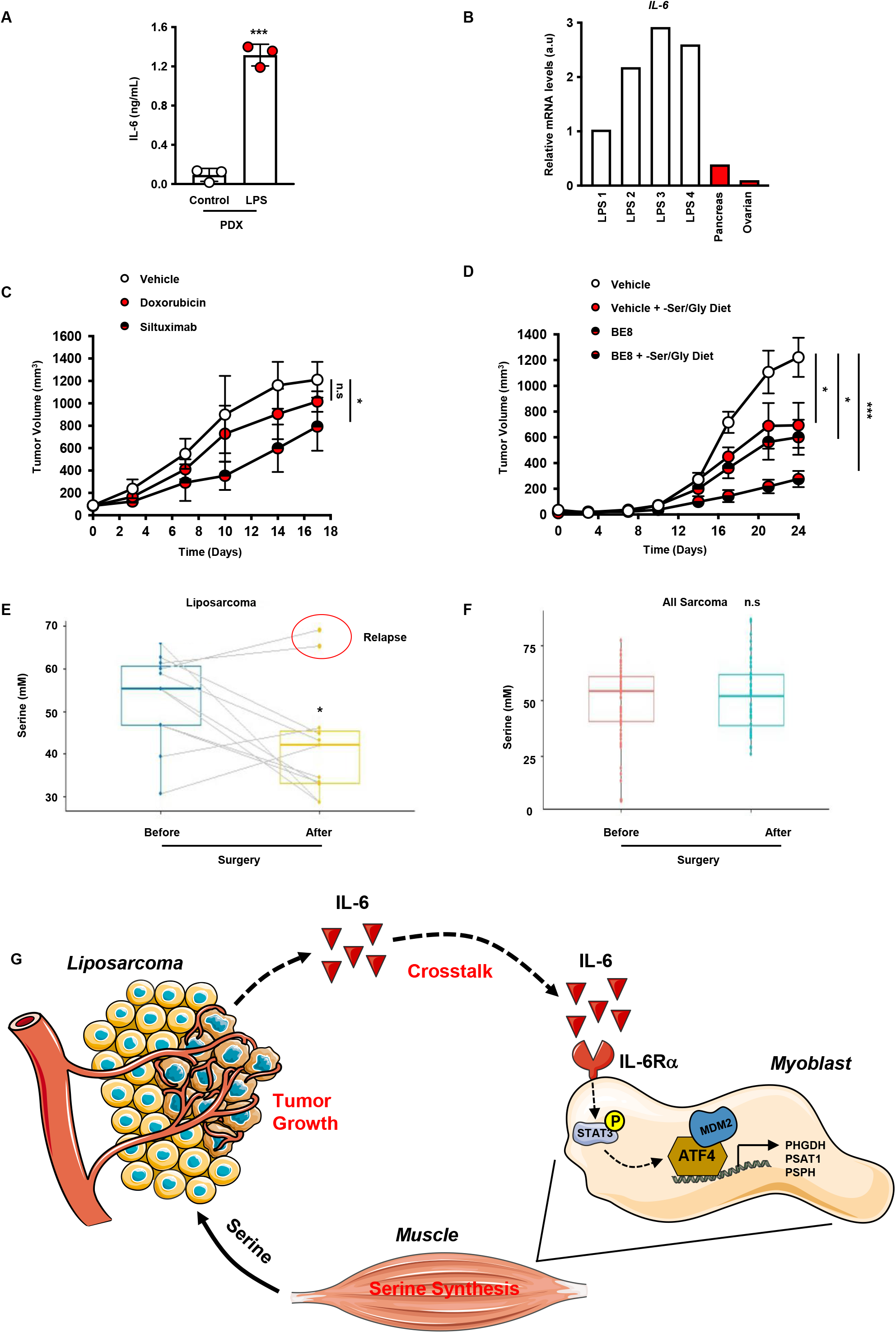
IL-6 inhibition as a new therapy in liposarcoma **A,** IL6 content (ng/ml), measured by Elisa. Control mice were compared to liposarcoma PDX Mice (LPS-PDX). **B,** Real-time qPCR analysis of *IL-6* mRNA level in frozen patient samples from liposarcoma, pancreas and ovarian cancer. **C,** Tumor growth curves from patient liposarcoma tumor subcutaneously implanted in nude mice treated or not with Siltuximab (10 mg/kg) or doxorubicine (2mg/kg) by IP weekly after tumor volume reached approximately 150mm^3^. Tumor volume was assessed at the indicated timepoints using caliper measurements (n=6 mice per group). **D,** Tumor growth curves from patient liposarcoma tumor subcutaneously implanted in nude mice, fed a normal or a no Serine/Glycine diet and treated or not with anti IL-6, BE8 (10 mg/kg) by IP weekly after tumor volume reached approximately 150mm^3^. Tumor volume was assessed at the indicated timepoints using caliper measurements (n=5 mice per group). **E** and **F,** Quantification of serum Serine levels (mM) from sarcoma patients before and 30 days after surgery, using liquid chromatography-high resolution mass spectrometry (LC/HRMS). **G,** Schematic representing crosstalk between liposarcoma tumors and surrounding muscles initiated through IL-6/STAT3 pathway activation. (All experiments were performed in at least triplicates and statistical analysis was applied with *=P<0.05, **=P<0.01, ***=<0.001, n.s=non-significant).

As well as guiding diagnostic, IL-6 could be exploited as a good target for treating liposarcoma. To assess the *in vivo* relevance of IL-6 pathway blockade, we combined Siltuximab, anti-IL-6 antibody, and doxorubicin, first-line treatment for primary LPS. PDX-LPS mice generated from patient samples were treated with Siltuximab (10 mg/kg twice a week) or doxorubicin (2 mg/kg, twice a week), and tumor growth was monitored. Anti-IL-6 antibody treatment decreased tumor growth, whereas doxorubicin had no statistically significant effect (Fig. 5C and Extended Fig. 5A). In contrast to doxorubicin, a 3-week treatment with Siltuximab did not result in obvious side effects in mice (data not shown). As validated by RT Q-PCR, *de novo* serine synthesis genes expression was decreased specifically in muscle tissue after Siltuximab treatment (Extended Fig. 5B, C).

Moreover, to evaluate the therapeutic use of our findings, suggesting that food-provided serine and IL-6-mediated blood serine elevation can promote liposarcoma tumor growth, we injected nude mice with LPS PDX and provided either a serine/glycine-free or normal diet. Then, mice were treated with anti-IL-6 leading to LPS tumor growth defects (Fig. 5D and Extended Fig. 5D). Interestingly, these data support the notion that serine/glycine metabolism initiated in muscle by LPS tumors is a driving event to sustain LPS tumor growth.

Collaboration between clinicians of the ‘Institut du Cancer de Montpellier’ (ICM), allowed us to generate a clinical-biobank (BCB) regrouping a collection of, fresh tumor and blood samples from LPS and other sarcoma patients. Using this biobank, we conducted a preliminary study on a small cohort and noticed that LPS patients exhibited higher serine blood levels before surgery and decreased drastically after surgery (Fig. 5E). Of note, serine levels did not move in other sarcoma subtypes (Fig. 5F). Two patients showed early recurrence and surprisingly, when looking at their blood serine concentration profile, we noticed that serine levels did not drop as low as levels observed in other patients (Fig. 5E). It seems essential to continue this study on a larger number of patients, to do so, clinical trial is ongoing on LPS patients to measure serine levels before and after surgery with a follow up of 2 years to anticipate potential recurrence.

Collectively, our findings reveal that liposarcoma cells reprogram distant muscle to meet their serine requirement and maintain proliferation through a STAT3/ATF4/C-MDM2 axis (Fig. 5F). Finally, serine levels could be use as early detection marker in liposarcoma and to predict recurrence after surgery. IL-6 may also be a new therapeutic strategy that, either singly or in combination, could benefit patients with liposarcoma.

## DISCUSSION

WD- and DD-LPS, the most frequent LPS subtypes, are poorly responsive to classical chemotherapies, and there is currently no cure available for metastatic or unresectable LPS. First line treatment for LPS is surgery and high doses of doxorubicin could be administrated, but it provides limited clinical benefit and commonly results in severe side effects (7). We recently demonstrated that in contrast to other types of sarcoma, all LPS display constitutive recruitment of C-MDM2 and are highly dependent on exogenous serine ^21^. In mammals, serine is considered a nonessential amino acid, the majority of which is synthesized *de novo* from glycolysis. Interestingly, many tumors such as triple-negative breast cancer and melanoma, up-regulate PHGDH expression leading to SSP flux increase and tumor growth *in vivo*. However, many tumors such as liposarcoma and lung cancer depend on the availability of extracellular serine suggesting that serine needs vary among cancer types. Nevertheless, liposarcoma cells consume large amounts of extracellular serine regardless of SSP activity and could affect exogenous serine availability. Counterintuitively, we observed that PDX mice with liposarcoma exhibit high level of circulating serine in blood compared to control mice fed with the same diet. We show here that surrounding muscles provide extracellular serine to sustain liposarcoma proliferation. Generating LPS-PDXs mice, which increased serum serine levels, demonstrate that *de novo* serine synthesis machinery was activated in these mice muscles. This was surprising, due to the distance between muscle and tumor, but muscle has already been described as a signaling organ releasing cytokines and metabolites during exercise. Our data reveal that conditioned media from LPS proliferating cells was sufficient to increase serine synthesis genes in muscle murine cell lines. The underlying signaling events linking LPS proliferation and muscle serine metabolism are likely to be complex but previous studies revealed that tumorkines are intimately involved in the initiation and progression of cancer, and circulating levels of many tumorkines are elevated in diverse cancer types. We found that IL-6 expression and concentration were elevated in LPS tumors compared to other cancer types tested. Using functional *in vitro* and *in vivo* studies, we demonstrate that LPS cells establish a metabolic cooperation with normal surrounding muscles to sustain serine requirements via IL-6 release. We observed that, using IL-6 pathways, tumor had the amazing power to reprogram distant muscle and understanding tumor biology beyond their own metabolic regulation seems to be essential to develop new therapeutic strategies. The molecular basis of increased IL-6 expression is mediated through MDM2/ATF4 chromatin function involved in serine homeostasis in LPS cells. As for MDM2, we provide genetic and pharmacological evidence that targeting IL-6-associated metabolic functions represents an efficient alternative strategy for LPS. IL-6 is a cytokine that plays roles in immunity, tissue regeneration, and metabolism. Production of IL-6 contributes to the defense during infection, but dysregulation of IL-6 signaling is also involved in disease pathology. The biological role of IL-6 has been implicated in various autoimmune disease such as rheumatoid arthritis and some other acute and chronic inflammation^37^. In addition to its critical role in several disease, IL-6 has very recently been reported key in the pathogenesis of multicentric Castleman disease (MCD)^38^. These features and our data make IL-6 and IL6-Rα attractive therapeutic targets, leading to the development of IL-6 pathway inhibitors including siltuximab, which has been granted full approval by the food and drug administration (FDA). In addition to our *in vitro* and *in vivo* experiments using siltuximab, further studies will be required to assess the efficacy and clinical utility of repurposing Siltuximab for liposarcoma therapy. Finally, our work highlights clearly the intriguing and emerging metabolic field of tumor - organ communication and may also suggest new combination therapies.

## Methods

All material and methods used are described in the Extended data.

## Acknowledgements

The authors would like to thank G. Gadea and C. Teyssier for input and critical reading of the manuscript. We thank the CRB-ICM (BB-033-0059) for the tumor samples supplied for this study. We thank all members of the animal, imaging, and histology facilities (Unité Mixte de Service, UMS3426 Montpellier BioCampus) for technical help.

## Author contributions

G.M, R.R and L.K.L., designed the studies, interpreted the data, and wrote the manuscript; G.M., J.P.R., B.R.M., A.A., S.J. and M.Y.C. performed *in vitro* the experiments; L.G, G.M, L.K.L contributed to the *in vivo* experiments. N.F. provided medical expertise.

## Data and materials availability

All the data used for this study are present in the paper or the Supplementary Materials.

Correspondence and requests for materials should be addressed to Laetitia K Linares.

## EXPERIMENTAL MODEL AND SUBJECT DETAILS

### Reagents

Recombinant Human IL-6 protein, Bazedoxifène, C188-9, Stattic, Phosphatase inhibitor cocktail 2, all unlabeled amino acids, formic acid (98% LC-MS grade), acetonitrile (>99.9% LC-MS grade), and methanol (>99.9% LC-MS grade) were purchased from Sigma-Aldrich. Protease inhibitor complete EDTA-free was purchased from Roche. The MDM2 inhibitor SP141 was synthesized based on the published structure^1^. Siltuximab (SYLVANT®) and Tocilizumab were purchased at the “ICM” pharmacy. Elsilimomab (BE8) et mAB 1339 (humanized aIl-6) were purchased by Evitria.

### Cell culture

Unless otherwise stated, cell culture reagents were purchased from Gibco (Invitrogen). Cells were kept at 37 °C in a humidified 5% CO2 incubator and maintained in DMEM Glutamax supplemented with 10% FBS (Hyclone, Thermo Scientific) and 1% Penicilline/Streptavidin. For Serine and Glycine starvation proliferation experiments, cells were cultured in MEM lacking amino acids supplemented with 1% Dialysed serum (Sigma-Merck, F0392), 1% Penycillin/Streptavidin, alanine (430μM), asparagine (50 μM), aspartic acid (20 μM), glutamic acid (80 μM), proline (200 μM) with (7AA medium) or without (5AA medium) serine (150 μM) and Glycine (300 μM).

### Construct and viral transduction

The following constructs were used for knockdown experiments:

For human protein: pLKO.1_Puro shMDM2 (Sigma-Aldrich mission clone #3377), pLKO.1_Puro shATF4 (Sigma-Aldrich mission clone #13577), pLKO.1_Puro shIL-6 (Sigma-Aldrich mission clones #59205, #59206, #59207).

For Murine protein: pLKO.1_Puro shMDM2 (Sigma-Aldrich mission clones #302276), pLKO.1_Puro shATF4 (Sigma-Aldrich mission clones #301646, #301731, #71724), pLKO.1_Puro shPHGDH (Sigma-Aldrich mission clones #41624, #41627, #41625), pLKO.1_Puro shPSAT1 (Sigma-Aldrich mission clones #120417, #120420, #120421), pLKO.1_Puro shPSPH (Sigma-Aldrich mission clones #81493, #81494, #81497), pLKO.1_Puro shIL6-Rα (Sigma-Aldrich mission clones #375089, #68293, #305257).

Lentivirus was prepared by co-transfection of 293T cells with shRNA of interest along with packaging plasmids pMD2.G (AddGene, cat. 12259), psPAX2 (AddGene, cat. 12260) and Lipofectamine 2000 transfection reagent (Invitrogen, cat. 15338). Lentivirus-containing media was collected from plates at 48h post-transfection, filtered using a 0.45μm filter, and stored at −80°C. For viral transduction, cells were incubated with lentivirus-containing medium and 8 μg/mL polybrene for 6h. Cells were allowed to recover for another 48h before selection with puromycin. All experiments were performed with cells that survived puromycin selection and displayed knockdown of MDM2, ATF4, PHGDH, PSAT1, IL6 and IL6Rα as assayed by western blot.

C2C12 cells were transfected using Lipofectamine 2000 to express Serine synthesis reporter gene : *Phgdh* (C2C12 Luc-*Phgdh*) or *Psat1* (C2C12 Luc-*Psat1*). Transduced myoblasts cells were selected with Hygromycine (2 μg/ml, Invitrogen) for 72 h.

### Serine and Glycine starvation Assay

C2C12 cells were conditioned to produce serine by seeding them at 8K/insert (Starsted, #833930040), 5h later inserts were transferred to a 6 well-plate containing 80K of IBGFP, and incubated for 72h. IB115-GFP were seeded in DMEM 10%FBS at 20K/well in 6 wells plate (Falcon #353046) after 5 hours plates and insert were washed 2 times with PBS and 5AA or 7AA medium were added to each well. LPS Cells proliferation was monitored in real time for 3 to 6 days using the Incucyte S3 Live-Cell Analysis system, whole-well module. The confluence value of each well was automatically monitored by the Incucyte system for 3-6 days and expressed as a value representing relative confluence area. Normalized Confluence was calculated by dividing the Confluence for each time point by the original Confluence. Effects of serine and Glycine deprivation on cell proliferation were confirmed by manual counting after trypan blue exclusion performed at the end of the experiments.

### Co-culture assay

C2C12 and LPS-cells (IB111, IB115), were seeded in 6-well plate with a ratio 1:2: 90K C2C12, 180K LPS-cells, in 2ml DMEM medium, for 48h. Similarly, cells (ratio 1:2) were cultivated separately using an insert. These experiments were also carried out using the shRNA-mediated depleted cells or with addition of drugs. Cells in culture alone or co-culture with LPS-cells were treated or not with a blocking anti-IL-6 antibody (Elsilimomab (BE8), mAB 1339 (humanized BE8), Siltuximab, 10 μg/ml), an anti-IL-6 receptor (Tocilizumab (20 μg/ml)) or inhibitor (Bazedoxifène, 100nM), or STAT3 inhibitors such as C188-9 and Stattic (10nM).

### Conditioned media (CM) assay

Myoblasts were seeded at 100K/well in a 6-well plate. About 30h after, medium was changed and 100-200μl of LPS-cells conditioned media (quantity corresponding to an IL-6 concentration finale of 150-300 pg/ml) or Fresh media was added to each well. Cells were collected after 16 hours. IL6 recombinant experiments were done using same experimental setting but by adding 50pg/ml of Human IL-6 recombinant proteins instead of CM on C2C12 cells. All myoblast metabolic reprogramming assay data were normalized to the corresponding control samples (mean +/-SD; n=3 independent experiments) and statistical significance was evaluated using non-parametric Mann Whitney U tests.

### RNA extraction and quantitative PCR

Total mRNAs were prepared using TriZol Reagent (Invitrogen). cDNAs were synthesized from 1µg of total RNA using SuperScript III Reverse Transcriptase (Invitrogen). Real-time quantitative PCRs were performed on a LightCycler 480 SW 1.5 apparatus (Roche) using the Platinum Taq DNA polymerase (Invitrogen) and a SYBR Green mix containing 3 mM MgCl2 and 30 μM of each dNTP using the following amplification procedure: 45 cycles of 95°C for 4 s, 60°C for 10 s, and 72°C for 15 s. The relative mRNA copy number was calculated using Ct value and normalized to at 2 or 3 housekeeping genes. Sequence of primers used for qPCR are listed in table S1.

### Immunoblotting

Protein extracts were subjected to SDS-PAGE and immunoblotted with the following primary antibodies: Mouse monoclonal: TBP (Santa Cruz, sc-56795), β-tubulin (Sigma-Aldrich, T6199), MDM2 (clones 4B11, and 2A10, Cell signaling and Millipore), STAT3 (Cell Signaling, 9139). Rabbit polyclonal: p-STAT3 (Cell signaling, 9145T), PHGDH (Cell Signaling, 13428), PSAT1 (Sigma, SAB2108040), PSPH (Santa Cruz, sc-365183), HSP90α (Cell Signalling, 8165), TBP (Cell Signalling, *8515)*. The proteins of interest were then detected either by incubation with horseradish peroxidase-conjugated anti-mouse and anti-rabbit IgG (Cell Signaling) secondary antibodies and revealed using the Pierce ECL Western Blotting Substrate or the SuperSignal West Femto Maximum Sensitivity Substrate (Thermo Fisher Scientific), or 488-, 680-fluorescent Antibodies (Li-Cor). Quantification of immunoblots was performed by densitometric analysis of the corresponding bands using ImageStudio and ImageJ software.

### Il-6 concentration Assay.

IL-6 concentration in the supernatant of the different cancer cell lines, and mice Serum was measured by an immunoassay (*Murine or* Human IL-6 Standard TMB ELISA Development Kit Catalog Number: 900-T50) following the manufacturer’s instructions. Cells supernatant were collected after 48h of culture, and centrifuge at 1 200 rpm. Mice serum was prepared by collecting 500μL to 1ml of blood from mice upon sacrifice in anticoagulant (Heparin 10µl/tube, 5000UI/ml). Mice blood was then centrifuged at 800g for 20min without the brakes, serum was then transferred to a vial, and store frozen prior analysis.

### XG-6 viability assay

The IL-6 dependent cell line XG-6 was cultured at a concentration of 1,5* 10^5^ cells/ml in RPMI medium in presence of recombinant IL-6 (control, 2ng/ml) or supernatant from two different cell lines (MCF7 breast cancer cells or IB115 liposarcoma cells) for 72hours. The quantity of XG-6 cells (AU) was measured by manual counting after trypan blue exclusion and CellTiter-Glo® Luminescent Cell Viability Assay (Promega, # G7570)

### ChIP experiment assay

Cells were collected and processed as previously described by Riscal et al. (2016). The Immunoprecipitation was performed using an anti-MDM2 (Santa Cruz, sc-813), anti-ATF4 (Cell signaling, 11815S) or a polyclonal rabbit IgG (Cell Signaling, 2729S) that serve as a control. Samples were than analyzed by qPCR using primers specific to IL-6 and TRIB3 (table S2).

### HPLC

Mice and patients serum samples was prepared directly upon collection as follow: Their blood were collected in anticoagulant (Heparin, EDTA coated collection tubes), then centrifuged at 800g for 20min, without the brakes. 30% acid salicylic was added to the samples and centrifuged again, supernatant was transferred to a vial, and store frozen prior analysis. The measurement was performed using an Agilent HPLC and data were analyzed using R studio.

### *In vivo* Experiments

Mice liposarcoma PDX models were established in collaboration with the surgical and pathology departments of the “Institut du Cancer de Montpellier” (ICM) by inserting a human tumor fragment of approximately 40 mm ^3^ subcutaneously on 8-week-old Nude mice (Charles River). Volumetric measurements of xenografted tumors were performed every 3 days by the same person using a manual caliper (volume = length × width^2^/2). All animals were euthanized when the first animal reached the ethical endpoint (volume = 1000 cm^3^ or ulceration). During our experiments mice were either fed with control diet (called Amino Acid diet; TD 99366 Harlan ENVIGO) or a test diet (Harlan Envigo, TD 130775: diet lacking serine and glycine) during 4 weeks. The diets had equal caloric value (3.9 kCal/g), an equal amount of total amino acids (179.6 g/kg) and are in a form of kibbles for mice. Total food intake was controlled to be identical in all experimental groups. Mice were housed in a pathogen-free barrier facility in accordance with the regional ethics committee for animal warfare (n°CEEA-LR-12067). Anti-IL-6 antibody (BE8 and Siltuximab), SP141 and doxorubicin were administered by IP injection at the dose of 10mg/kg twice a week for 3 weeks, 40mg/kg weekly and 2mg/kg weekly respectively.

---

**Supplementary Figure 1:**
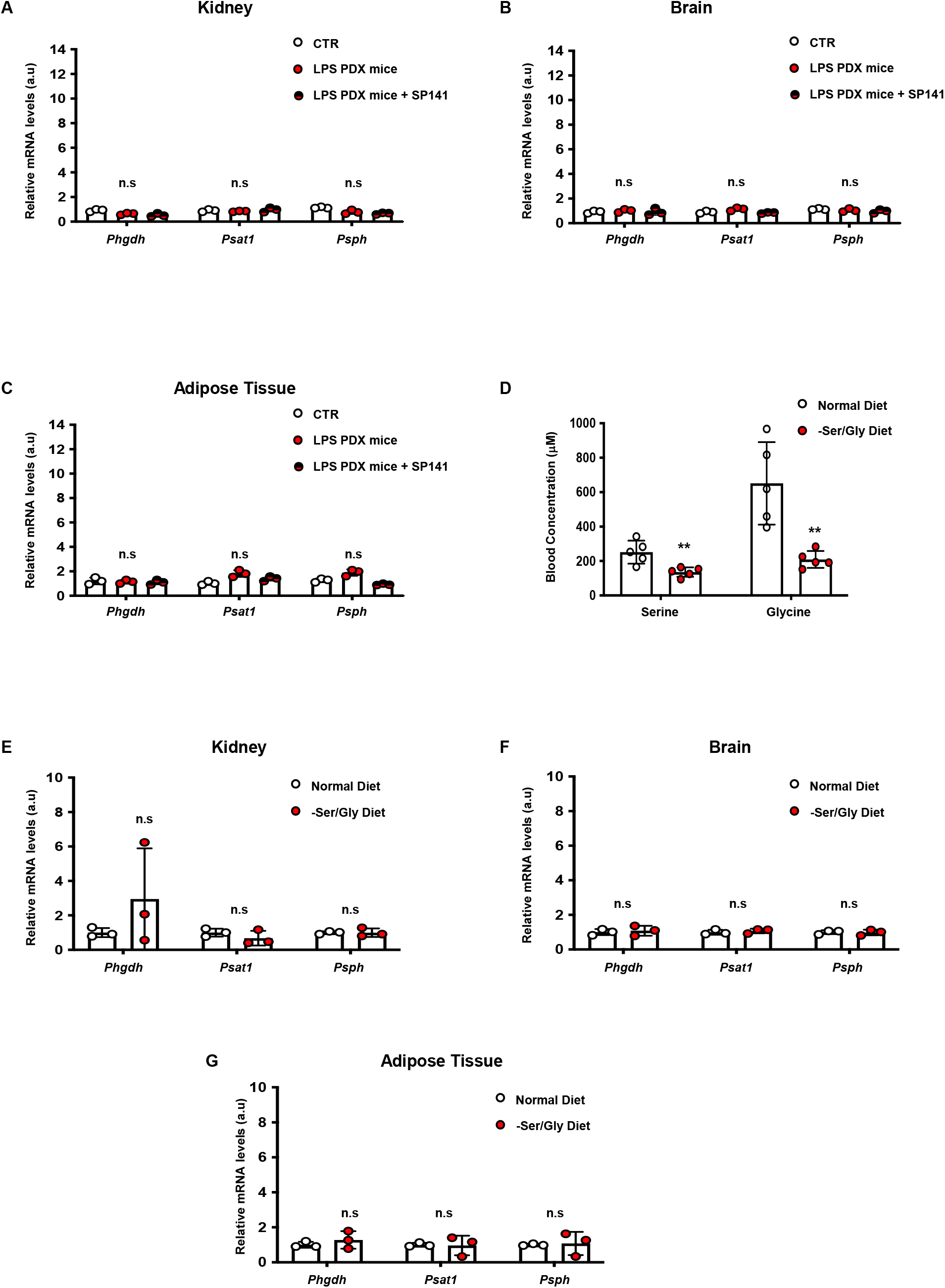
**A,** Real-time qPCR analysis performed on LPS-PDX and control mice kidney, evaluating expression of serine synthesis pathway genes: *Phgdh, Psat1* and *Psph*. Mice were treated daily with placebo or SP141 (40mg/ml). **B,** Real-time qPCR analysis performed on LPS-PDX and control mice brain, evaluating expression of serine synthesis pathway genes: *Phgdh, Psat1* and *Psph*. Mice were treated daily with placebo or SP141 (40mg/ml). **C,** Real-time qPCR analysis performed on LPS-PDX and control mice adipose tissue, evaluating expression of serine synthesis pathway genes: *Phgdh, Psat1* and *Psph*. Mice were treated daily with placebo or SP141 (40mg/ml). **D,** Serum Serine and Glycine levels (μM) measured by HPLC. Mice were fed with normal or -Ser/Gly diet. **E,** Real-time qPCR analysis performed on control mice kidney, evaluating expression of serine synthesis pathway genes: *Phgdh, Psat1* and *Psph*. Mice were fed with normal or -Ser/Gly diet. **F,** Real-time qPCR analysis performed on control mice brain, evaluating expression of serine synthesis pathway genes: *Phgdh, Psat1* and *Psph*. Mice were fed with normal or -Ser/Gly diet. **G,** Real-time qPCR analysis performed on control mice adipose tissue, evaluating expression of serine synthesis pathway genes: *Phgdh, Psat1* and *Psph*. Mice were fed with normal or -Ser/Gly diet. (All experiments were performed in at least triplicates and statistical analysis was applied with *=P<0.05, **=P<0.01, ***=<0.001, n.s=non-significant).

**Supplementary Figure 2:**
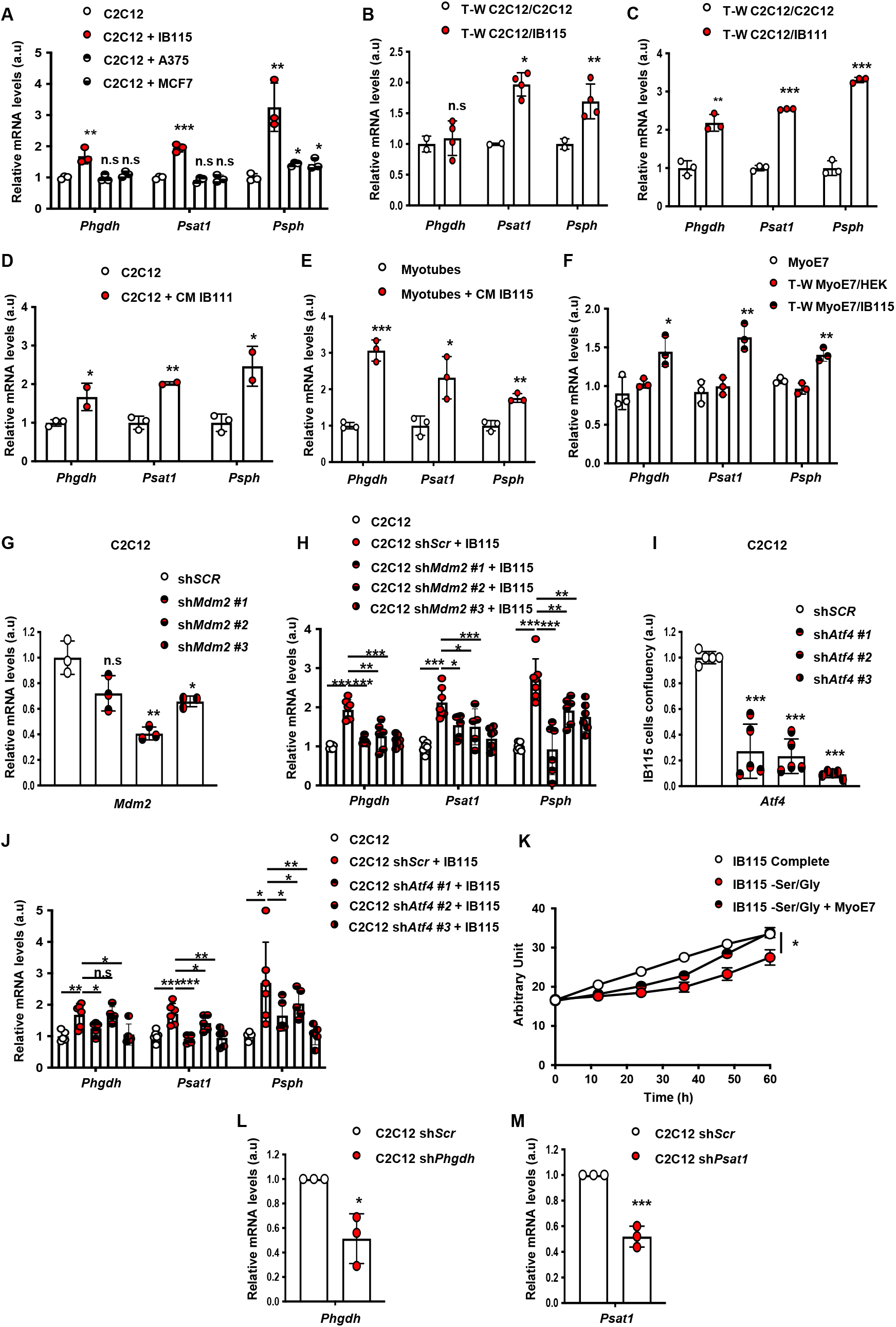
**A,** Real-time qPCR analysis performed on C2C12 cocultured with IB115, A375 and MCF7 cells (ratio 1:2), evaluating expression of serine synthesis pathway genes: *Phgdh, Psat1* and *Psph*. **B,** Real-time qPCR analysis performed on C2C12 cocultured in transwell for 48h with IB115 cells, evaluating expression of serine synthesis pathway genes: *Phgdh, Psat1* and *Psph*. **C,** Real-time qPCR analysis performed on C2C12 cocultured in transwell for 48h with IB111 cells, evaluating expression of serine synthesis pathway genes: *Phgdh, Psat1* and *Psph*. **D,** Real-time qPCR analysis performed on C2C12 differentiated in myotubes incubated 16h with IB115 conditioned media, evaluating expression of serine synthesis pathway genes: *Phgdh, Psat1* and *Psph*. **E,** Luciferase assay performed on C2C12 cells incubated 16h with IB115 conditioned media, evaluating relative luciferase activity of *Phgdh* and *Psat1* reporter. **F,** Real-time qPCR analysis performed on human myoblasts MyoE7 cocultured in transwell for 48h with IB115 or HEK cells, evaluating expression of serine synthesis pathway genes: *Phgdh, Psat1* and *Psph*. **G,** Real-time qPCR analysis of *Mdm2* mRNA level in C2C12 cells after sh*Scr* or 3 different sh*Mdm2* lentiviral infection, puromycin selection (48h, 2μg/mL).**H,** Real-time qPCR analysis performed on C2C12 cells after sh*Scr* or 3 different sh*Mdm2* cocultured with IB115 cells (ratio 1:2), evaluating expression of serine synthesis pathway genes: *Phgdh, Psat1* and *Psph*. **I,** Real-time qPCR analysis of *Atf4* mRNA level in C2C12 cells after sh*Scr* or 3 different sh*Atf4* lentiviral infection, puromycin selection (48h, 2mg/mL). **J,** Real-time qPCR analysis performed on C2C12 cells after sh*Scr* or 3 different sh*Atf4* cocultured with IB115 cells (ratio 1:2), evaluating expression of serine synthesis pathway genes: *Phgdh, Psat1* and *Psph*. **K,** Proliferation assay performed on IB115 cells grown in media supplemented with or without Serine and Glycine and cocultured with human myoblasts MyoE7. **L,** Real-time qPCR analysis of *Phgdh* mRNA level in C2C12 cells after sh*Scr* or sh*Phgdh* lentiviral infection, puromycin selection (48h, 2mg/mL). **M,** Real-time qPCR analysis of *Psat1* mRNA level in C2C12 cells after sh*Scr* or sh*Psat1* lentiviral infection, puromycin selection (48h, 2mg/mL). (All experiments were performed in at least triplicates and statistical analysis was applied with *=P<0.05, **=P<0.01, ***=<0.001, n.s=non-significant).

**Supplementary Figure 3:**
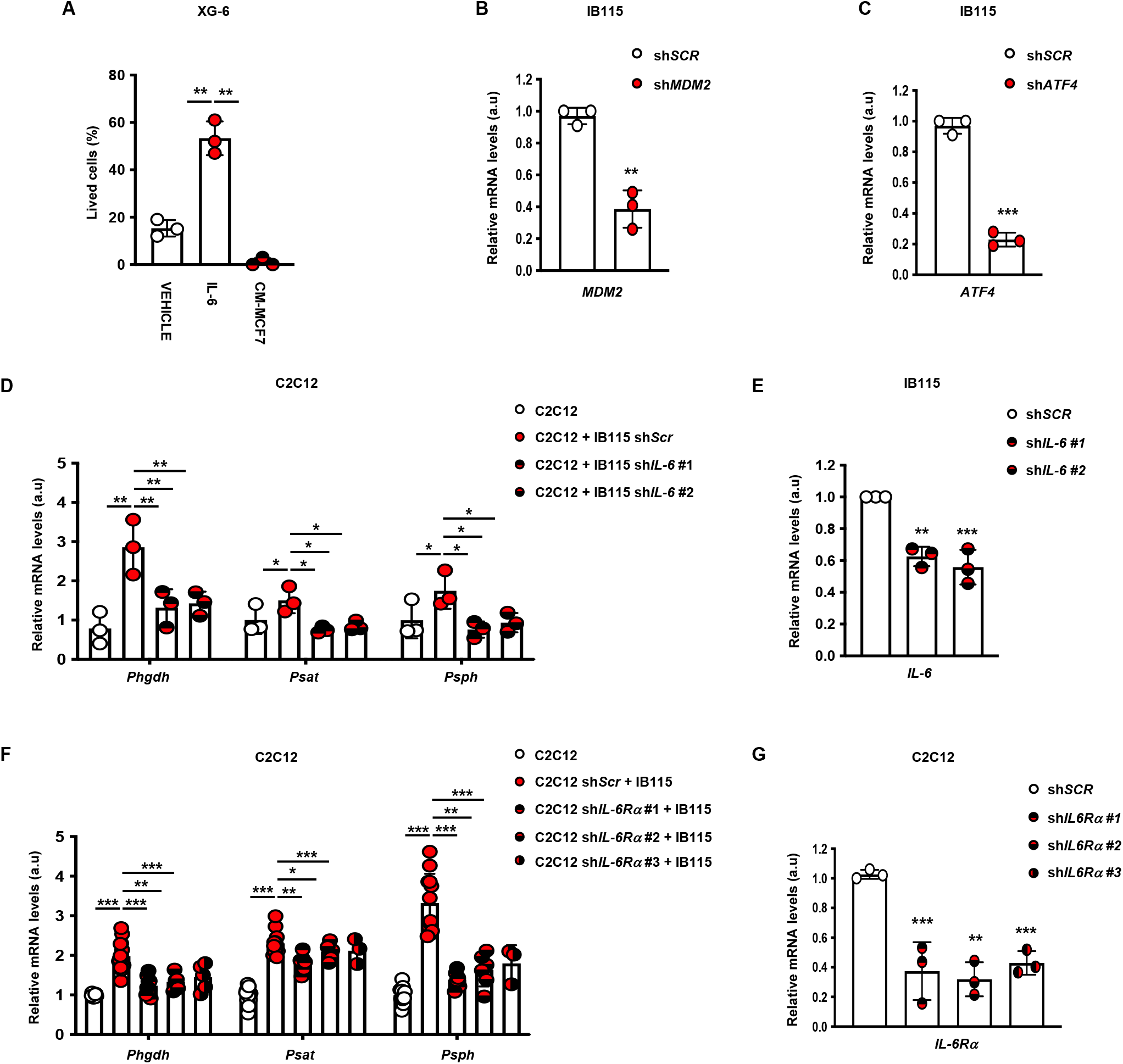
**A,** End point of proliferation assay performed on XG-6 cells grown in media supplemented with vehicle, recombinant IL-6 or MCF7 conditioned media. **B,** Real-time qPCR analysis of *MDM2* mRNA level in IB115 cells after sh*SCR* or sh*MDM2* lentiviral infection, puromycin selection (48h, 2μg/mL). **C,** Real-time qPCR analysis of *ATF4* mRNA level in IB115 cells after sh*SCR* or sh*ATF4* lentiviral infection, puromycin selection (48h, 2μg/mL). **D,** Real-time qPCR analysis performed on C2C12 cells cocultured with IB115 cells after sh*SCR* or 2 different sh*IL-6* (ratio 1:2), evaluating expression of serine synthesis pathway genes: *Phgdh, Psat1* and *Psph*. **E,** Real-time qPCR analysis of *IL-6* mRNA level in IB115 cells after sh*Scr* or 2 different sh*IL-6* lentiviral infection, puromycin selection (48h, 2μg/mL). **F,** Real-time qPCR analysis performed on C2C12 cells after sh*SCR* or 3 different sh*IL-6Rα* cocultured with IB115 cells (ratio 1:2), evaluating expression of serine synthesis pathway genes: *Phgdh, Psat1* and *Psph*. **G,** Real-time qPCR analysis of *IL-6Rα* mRNA level in C2C12 cells after sh*Scr* or 3 different sh*IL-6Rα* lentiviral infection, puromycin selection (48h, 2μg/mL). (All experiments were performed in at least triplicates and statistical analysis was applied with *=P<0.05, **=P<0.01, ***=<0.001, n.s=non-significant).

**Supplementary Figure 4:**
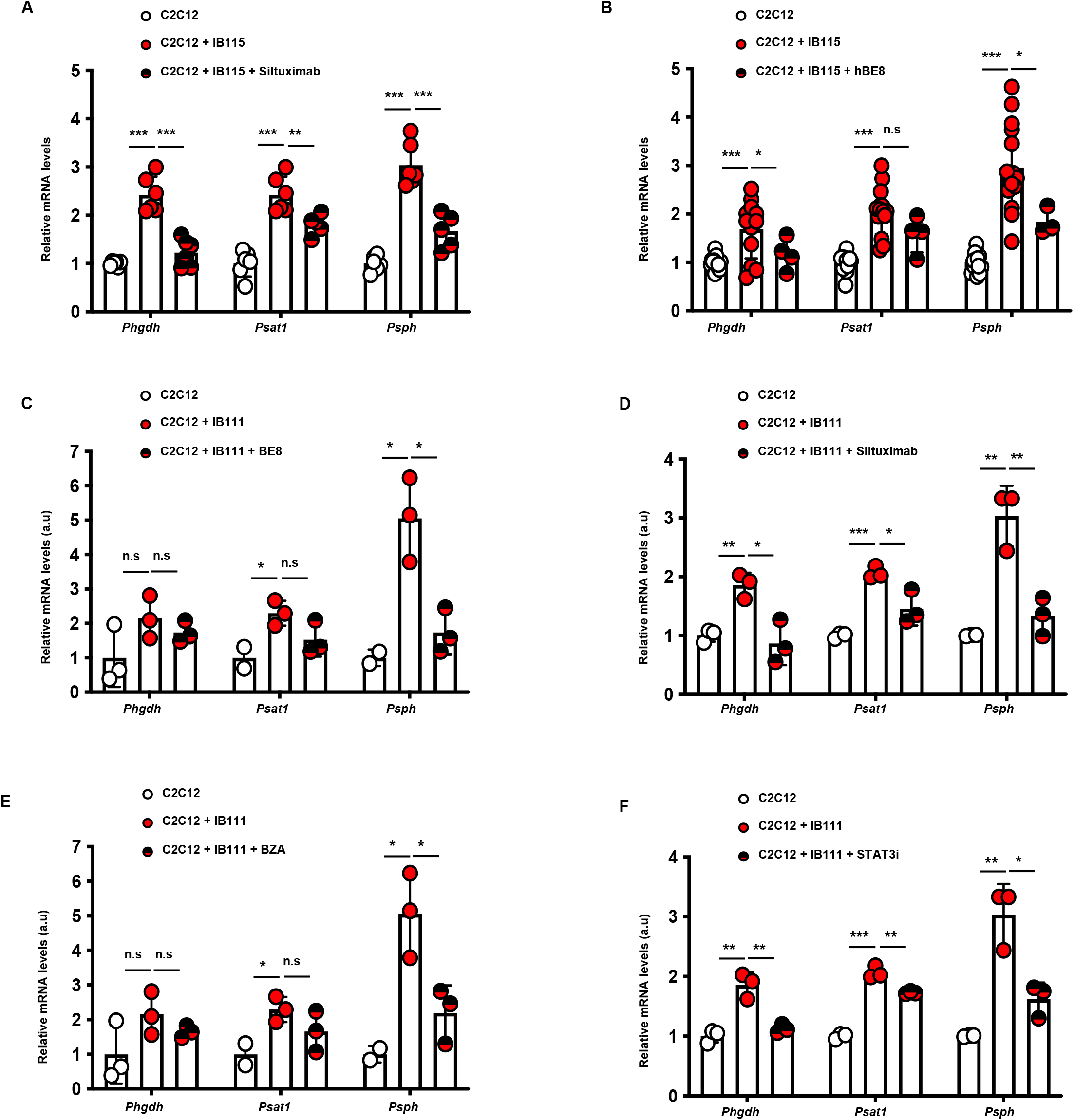
**A,** Real-time qPCR analysis performed on C2C12 cells alone or cocultured with IB115 cells grown in media supplemented with vehicle or siltuximab, evaluating expression of serine synthesis pathway genes: *Phgdh, Psat1* and *Psph*. **B,** Real-time qPCR analysis performed on C2C12 cells alone or cocultured with IB115 cells grown in media supplemented with vehicle or anti IL-6 (hBE8), evaluating expression of serine synthesis pathway genes: *Phgdh, Psat1* and *Psph*. **C,** Real-time qPCR analysis performed on C2C12 cells alone or cocultured with IB111 cells grown in media supplemented with vehicle or anti IL-6 (BE8), evaluating expression of serine synthesis pathway genes: *Phgdh, Psat1* and *Psph*. **D,** Real-time qPCR analysis performed on C2C12 cells alone or cocultured with IB111 cells grown in media supplemented with vehicle or siltuximab, evaluating expression of serine synthesis pathway genes: *Phgdh, Psat1* and *Psph*. **E,** Real-time qPCR analysis performed on C2C12 cells alone or cocultured with IB111 cells grown in media supplemented with vehicle or GP130 inhibitor (BZA), evaluating expression of serine synthesis pathway genes: *Phgdh, Psat1* and *Psph*. **F,** Real-time qPCR analysis performed on C2C12 cells alone or cocultured with IB111 cells grown in media supplemented with vehicle or STAT3 inhibitor, evaluating expression of serine synthesis pathway genes: *Phgdh, Psat1* and *Psph*. (All experiments were performed in at least triplicates and statistical analysis was applied with *=P<0.05, **=P<0.01, ***=<0.001, n.s=non-significant).

**Supplementary Figure 5:**
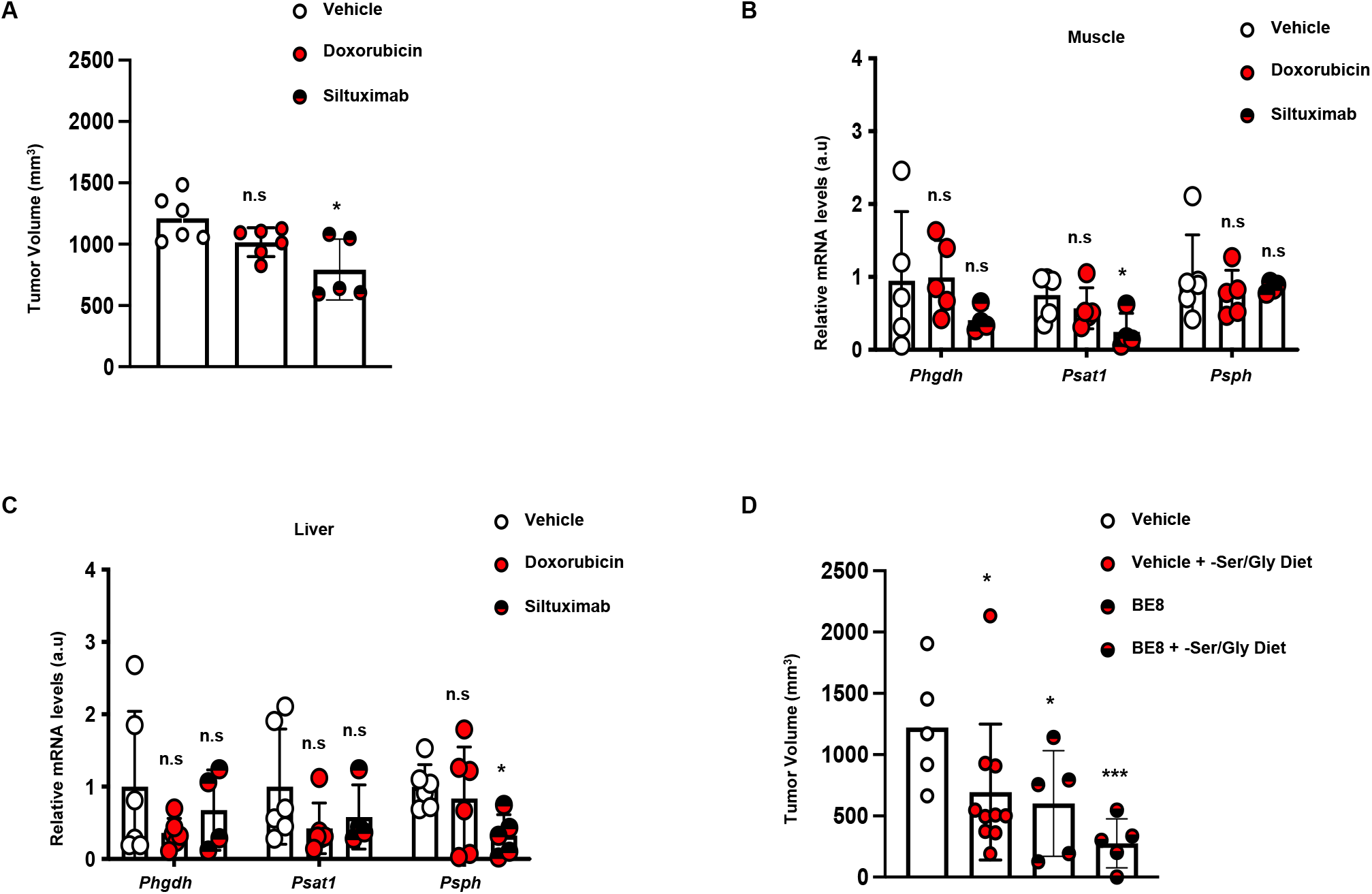
**A,** Tumor weight from patient liposarcoma tumor subcutaneously implanted in nude mice treated or not with Siltuximab (10 mg/kg) or doxorubicine (2mg/kg), 17 days after implantation. **B,** Real-time qPCR analysis performed on mice muscle, evaluating expression of serine synthesis pathway genes: *Phgdh, Psat1* and *Psph*. Mice were treated or not with Siltuximab (10 mg/kg) or doxorubicine (2mg/kg) by IP weekly **C,** Real-time qPCR analysis performed on mice liver, evaluating expression of serine synthesis pathway genes: *Phgdh, Psat1* and *Psph*. Mice were treated or not with Siltuximab (10 mg/kg) or doxorubicine (2mg/kg) by IP weekly **D,** Tumor weight from patient liposarcoma tumor subcutaneously implanted in nude mice, fed a normal or a no Serine/Glycine diet and treated or not with anti IL-6, BE8 (10 mg/kg), 24 days after implantation. (All experiments were performed in at least triplicates and statistical analysis was applied with *=P<0.05, **=P<0.01, ***=<0.001, n.s=non-significant).

